# Cross-Modal Benchmarking of Acoustic Prosody and Ventral Striatal BOLD for Depression-Related Anhedonia Classification: A Pre-Registered Study with the ClinicalWhisper Pipeline

**DOI:** 10.64898/2026.06.08.728970

**Authors:** Chengdong Zhou, Mei-Hui Wu, Yi-Ming Xiang, Laurent Itti

## Abstract

Can computational analysis of a brief voice recording classify depression-related anhedonia as effectively as task-based fMRI? We address this question through a pre-registered cross-modal benchmarking study (osf.io/bsvrj) that evaluates two independent classification pipelines against depression-related anhedonia operationalized via self-report. Anhedonia— the diminished capacity to experience pleasure or motivation to pursue rewards—is a transdiagnostic marker of reward-system dysfunction that predicts treatment resistance in major depressive disorder, yet current assessment requires either expensive functional neuroimaging or subjective self-report scales, neither of which scales to routine screening. We benchmarked two independent classification pipelines: **Stream A** extracted 88 acoustic prosody features from DAIC-WOZ clinical interviews (*n* = 142) using the ClinicalWhisper pipeline (Whisper Large-v3, pyannote diarization, OpenSMILE eGeMAPS v02); **Stream B** extracted nucleus accumbens BOLD activation during a reward task from the UCLA ds000030 dataset (*n* = 272). Three classifiers (logistic regression, random forest, gradient-boosted trees) were evaluated under stratified 5-fold cross-validation with fixed, pre-registered hyperparameters.

Stream A achieved a best AUC-ROC of 0.63 (random forest; permutation *p* = .049), confirming H1 that acoustic prosody classifies PHQ-8-defined anhedonia above chance at *α* = .05 (uncorrected), though this result does not survive Bonferroni correction for three classifiers (*α/*3 = .017). Bootstrap analysis of ΔAUC confirmed non-inferiority relative to Stream B (H2: 95% CI lower bound *> −*0.10). However, Stream B itself did not achieve above-chance classification (*p* = .057), so this non-inferiority finding reflects comparable, modest performance across both modalities rather than equivalence to a validated neural biomarker. Pitch variability features (F0) ranked among the top 5 predictors by SHAP value in the gradient-boosted trees model (H3: partially supported). An exploratory combined model (eGeMAPS + recurrence quantification analysis) reached AUC = 0.65 (GBT; *p* = .032), though this lift was not statistically significant by DeLong test (*z* = *−*0.44, *p* = .66).

These results provide initial evidence that vocal prosody, acquired through a standardized, open-source pipeline, carries depression-related information comparable to fMRI-derived ventral striatal activation for binary classification. However, neither stream’s operationalization isolates anhedonia from general depressive symptomatology, and the cross-modal design compares different constructs across different cohorts. We refer to the classification target throughout as the “anhedonia composite” to acknowledge that PHQ-8 Items 1+2 conflate anhedonia with dysphoria. We discuss these constraints and their implications for the causal framework motivating this work.

## 1 Introduction

Major depressive disorder (MDD) is the leading cause of disability worldwide, affecting an estimated 280 million people and accounting for more years lived with disability than any other condition (World Health Organization, 2023). Global prevalence increased by approximately 25% during the first year of the COVID-19 pandemic alone (Santomauro et al., 2021), underscoring both the scale and the trajectory of the crisis. Despite widespread availability of first-line pharmacotherapies, approximately one-third of patients fail to achieve adequate remission (Rush et al., 2006), and treatment-resistant presentations disproportionately involve anhedonia, defined as the diminished capacity to experience pleasure or motivation to pursue rewards (Spijker et al., 2001). First included among diagnostic criteria for depression in the Feighner criteria (Feighner et al., 1972) and retained in every subsequent edition of the DSM, anhedonia is now recognized not merely as a symptom but as a transdiagnostic marker of reward-system dysfunction that cuts across MDD, schizophrenia, substance use disorders, and Parkinson’s disease (Husain & Roiser, 2018; Treadway & Zald, 2011).

Contemporary neuroscience has localized the neural substrate of anhedonia to dopaminergic circuits projecting from the ventral tegmental area (VTA) to the nucleus accumbens (NAcc) and medial prefrontal cortex (Pizzagalli et al., 2009; Treadway et al., 2012). Patients with elevated anhedonia consistently show blunted NAcc BOLD activation during reward anticipation in fMRI paradigms, most cleanly demonstrated with the Monetary Incentive Delay task (Knutson et al., 2000). Berridge and Robinson’s influential framework distinguishes “wanting” (incentive salience, driven by mesolimbic dopamine) from “liking” (hedonic impact, driven by opioidergic hotspots in the NAcc shell), and this dissociation has received substantial support from both animal models and human imaging (Berridge & Robinson, 1998; Treadway et al., 2012). In principle, this distinction suggests that behavioral and physiological signals downstream of dopaminergic circuitry may carry information about motivational anhedonia specifically, though isolating this component from general dysphoria in clinical populations remains a fundamental measurement challenge.

Vocal prosody is one candidate downstream signal. Speech production recruits laryngeal and respiratory motor control circuits, and there is evidence from Parkinson’s disease that dopaminergic depletion produces characteristic vocal changes: reduced pitch variability, compressed loudness range, and imprecise articulation (Harel et al., 2004; Skodda et al., 2011). Simonyan, Horwitz, and Jarvis reviewed the comparative neurobiology of dopamine’s role in vocal production across species and concluded that dopaminergic modulation of vocal motor circuits is phylogenetically conserved, though the specific pathway from mesolimbic dopamine to laryngeal motor nuclei in humans remains incompletely mapped (Simonyan et al., 2012). Depressive speech shows parallel prosodic flattening, including reduced pitch variability, compressed dynamic range, and slower speaking rate (Cummins et al., 2015). Yet the existing vocal-biomarker literature has focused almost exclusively on global depression severity (e.g., PHQ-8 total score) rather than anhedonia specifically (Low et al., 2020; Scherer et al., 2014). Furthermore, most prior studies employ proprietary, non-reproducible feature extraction pipelines, limiting clinical translation and cross-study comparison (Cummins et al., 2015). Prior entries in the Audio/Visual Emotion Challenge (AVEC) depression sub-challenges achieved AUCs in the range of 0.60–0.75 for depression severity classification on the same DAIC-WOZ corpus (Valstar et al., 2016), providing a benchmark against which the present results should be interpreted.

The present study asks whether a five-minute voice recording can classify anhedonia (operationalized via self-report) at a level comparable to fMRI-derived ventral striatal activation. We introduce ClinicalWhisper, an open-source, standardized acoustic processing pipeline (Whisper Large-v3, pyannote 3.1 speaker diarization, OpenSMILE eGeMAPS v02) designed for reproducible clinical speech analysis. We pre-registered three hypotheses (osf.io/bsvrj): (H1) acoustic prosody classifies anhedonia above chance; (H2) acoustic classification is non-inferior to fMRI-based classification (ΔAUC lower CI *> −*0.10); and (H3) pitch and energy features rank among the top predictors.

Two design constraints should be stated at the outset. First, because no publicly available dataset pairs naturalistic clinical audio with task-based fMRI and validated anhedonia labels, the two streams draw from separate cohorts (DAIC-WOZ and UCLA ds000030), making this a cross-modal benchmarking study rather than a multimodal fusion design. Second, the two streams use different anhedonia operationalizations: PHQ-8 Items 1+2 for Stream A and the Chapman Physical Anhedonia Scale for Stream B. These measure overlapping but distinct constructs, and neither cleanly isolates motivational anhedonia from dysphoria or consummatory deficits. We discuss the implications of this construct heterogeneity directly.

## 2 Methods

The authors have observed the data but have not yet performed the proposed analyses at the time of pre-registration. All analyses reported here were conducted after the pre-registration was time-stamped.

### 2.1 Datasets

This study employed a cross-modal benchmarking design comparing two independent classification streams. Because no publicly available dataset includes both naturalistic clinical audio and task-based fMRI with validated anhedonia labels, each stream draws from a separate corpus. This design was pre-registered prior to analysis (osf.io/bsvrj).

#### 2.1.1 DAIC-WOZ (Stream A: Acoustic Prosody)

The Distress Analysis Interview Corpus, Wizard of Oz edition (DAIC-WOZ; Gratch et al. 2014) comprises 189 semi-structured clinical interviews conducted by a virtual interviewer, with accompanying audio recordings and Patient Health Questionnaire (PHQ-8) responses. DAIC-WOZ is the standard evaluation corpus for the AVEC depression analysis challenge series (Valstar et al., 2016).

Anhedonia severity was operationalized as the sum of PHQ-8 Items 1 (“little interest or pleasure in doing things”) and 2 (“feeling down, depressed, or hopeless”), yielding a subscale range of 0–6. Participants with a summed score *≥* 3 were classified as *anhedonic*; all others as *non-anhedonic*. This operationalization follows the pre-registration and is consistent with prior AVEC challenge work, but it carries an important limitation: Item 2 captures dysphoria (depressed mood), not anhedonia. The composite therefore conflates anhedonia with general depressive symptomatology. We return to this construct validity constraint in the Discussion.

The *≥* 3 threshold was selected as part of the pre-registration to approximate moderate-to-severe endorsement of either item, though no published psychometric work validates this specific cutoff as a clinically meaningful anhedonia boundary on the PHQ-8 Items 1+2 subscale. After exclusions for failed transcription (*n* = 12), failed diarization (*n* = 8), missing PHQ-8 (*n* = 3), and audio quality (SNR *<* 10 dB; *n* = 24), the final analytic sample comprised *n* = 142 participants (32 anhedonic, 110 non-anhedonic; 22.5% prevalence).

#### 2.1.2 UCLA ds000030 (Stream B: Neural Activation)

The UCLA Consortium for Neuropsychiatric Phenomics dataset (ds000030; Poldrack et al. 2016) provides task-based fMRI data from 272 participants spanning healthy controls and individuals with schizophrenia, bipolar disorder, or ADHD. The Balloon Analogue Risk Task (BART; Lejuez et al. 2002) was administered during fMRI acquisition. After exclusions for excessive motion, failed normalization, and missing Chapman PAS data, the final analytic sample comprised *n* = 234 participants.

A methodological note on task choice: the BART is a reward-processing paradigm, but unlike the Monetary Incentive Delay (MID) task, which was designed specifically to separate reward anticipation from reward receipt (Knutson et al., 2000), the BART conflates reward processing with risk-taking and impulsivity. The *cash-out > explode* contrast captures risk-related decision outcomes rather than the anticipatory reward signal most directly linked to NAcc dopamine. This constrains the specificity of Stream B as a “reward-circuit” benchmark: elevated BART activation may reflect risk tolerance or impulsivity rather than reward anticipation per se.

Anhedonia was operationalized using the Revised Chapman Physical Anhedonia Scale (RPAS; Chapman & Chapman 1978), a 61-item true–false questionnaire assessing diminished sensory and consummatory pleasure. RPAS total scores were binarized at the sample median, classifying participants into high-anhedonia and low-anhedonia groups. This median-split approach is statistically problematic: dichotomizing a continuous variable at an arbitrary point discards variance, misclassifies individuals near the median, and inflates Type II error (MacCallum et al., 2002). Furthermore, the RPAS measures a fundamentally different anhedonia construct than the PHQ-8 Items 1+2 composite used in Stream A. The RPAS assesses consummatory/sensory pleasure deficits (“I have always found organ music dull and unexciting”), whereas PHQ-8 Item 1 captures interest/pleasure more broadly and Item 2 captures dysphoria. The “wanting” versus “liking” dissociation that motivates our theoretical framework (Berridge & Robinson, 1998) cannot be resolved by either measure. This construct heterogeneity means that the cross-modal comparison is not strictly apples-to-apples at the construct level, a limitation that bounds the interpretive scope of H2.

#### 2.1.3 Ethics Statement

Both datasets consist of pre-existing, de-identified data. DAIC-WOZ access was obtained under a signed data use agreement with the USC Institute for Creative Technologies. ds000030 is openly available on OpenNeuro under the PDDL license. This study was submitted for review by the USC Institutional Review Board (Protocol UP-26-00343); a determination is pending. Because both datasets are pre-existing and fully de-identified, this work is expected to qualify as not human subjects research under 45 CFR 46.104(d)(4).

### 2.2 Stream A: ClinicalWhisper Acoustic Processing Pipeline

All acoustic processing was performed using ClinicalWhisper, an open-source pipeline developed for this study and publicly available at https://github.com/czhou732/ClinicalWhisper. The pipeline integrates three components in sequence:

#### Automatic Speech Recognition (ASR)

Audio recordings were transcribed using OpenAI Whisper Large-v3 (Radford et al., 2023), a 1.5-billion-parameter transformer model trained on 680,000 hours of multilingual speech. Whisper was selected for its state-of-the-art word error rate on clinical speech and robustness to background noise. Processing was performed locally on USC igpu cluster GPU nodes to maintain HIPAA-compliant data handling (no audio transmitted to external servers).

#### Speaker Diarization

Interviewer and participant speech segments were separated using pyannote.audio 3.1 (Bredin, 2023), a neural speaker diarization pipeline based on temporal convolutional networks. Only participant speech segments were retained for feature extraction, eliminating interviewer-induced variance. Diarization quality was verified by manual inspection of a 10% random sample (*n* = 19); segment-level agreement exceeded 95%.

#### Acoustic Feature Extraction

The extended Geneva Minimalistic Acoustic Parameter Set (eGeMAPS v02; Eyben et al. 2016) was extracted from participant speech segments using OpenSMILE. eGeMAPS v02 comprises 88 low-level descriptors spanning five acoustic domains: prosodic features (fundamental frequency [F0] mean, coefficient of variation, percentiles, rising/falling slopes), energy features (loudness mean, coefficient of variation, percentiles, spectral flux), spectral features (MFCCs 1–4, alpha ratio, Hammarberg index, spectral slopes), voice quality features (jitter, shimmer, harmonics-to-noise ratio [HNR]), and temporal features (voiced/unvoiced segment lengths, speaking rate, pause duration).

Features were aggregated at the participant level using five summary statistics across all speech segments: mean, standard deviation, and the 10th, 50th, and 90th percentiles, yielding a final feature matrix of 447 features per participant after removing zero-variance columns from the 89 base features *×* 5 aggregation statistics.

#### Recurrence Quantification Analysis (RQA; Post-Hoc Exploratory)

As a post-hoc exploratory analysis (not included in the pre-registration), recurrence quantification analysis was applied to the time series of each eGeMAPS feature to capture temporal dynamics not represented in static summary statistics. RQA features (recurrence rate, determinism, laminarity, entropy, trapping time) were computed for each of 15 base acoustic features, yielding 74 additional features per participant. Combined eGeMAPS + RQA feature sets totaled 521 features.

### 2.3 Stream B: fMRI Processing Pipeline

#### Preprocessing

Structural and functional images were preprocessed using fMRIPrep 23.x (Esteban et al., 2019), which performs brain extraction, spatial normalization to MNI152NLin2009cAsym template space (2 mm isotropic), motion correction, susceptibility distortion correction, and confound estimation. Spatial smoothing was applied at 6 mm FWHM Gaussian kernel.

#### Quality Control

Participants were excluded for: mean framewise displacement (FD) *>* 0.5 mm, *>* 20% of frames censored, failed spatial normalization (visual inspection), or missing Chapman PAS data. Motion regressors (six rigid-body parameters and their temporal derivatives) were included as nuisance covariates in the first-level GLM.

#### First-Level GLM

A subject-level general linear model was estimated using Nilearn (Abraham et al., 2014) with an AR(1) noise model. The BART task was modeled with regressors for cash-out trials and explode trials, convolved with the canonical hemodynamic response function. The contrast of interest was *cash-out > explode*, isolating reward-related neural activation.

#### ROI Extraction

Mean BOLD percent signal change was extracted from bilateral nucleus accumbens, caudate, and putamen using the Harvard-Oxford subcortical atlas (Jenkinson et al., 2012). The primary feature for classification was bilateral NAcc activation; caudate and putamen were included as secondary features to capture broader dorsal striatal involvement.

### 2.4 Classification Framework

Identical classification procedures were applied to both streams, as pre-registered.

#### Classifiers

Three classifiers spanning complementary model families were trained: L2-regularized logistic regression (*C* = 1.0), random forest (*n*_estimators_ = 500, max_depth = None), and gradient-boosted trees (scikit-learn GradientBoostingClassifier; *n*_estimators_ = 300, learning rate = 0.1).

Hyperparameters were fixed a priori per the pre-registration; no nested tuning was performed. All classifiers were implemented in scikit-learn (Pedregosa et al., 2011).

#### Cross-Validation

Stratified 5-fold cross-validation preserved class proportions in each fold. The primary evaluation metric was AUC-ROC (area under the receiver operating characteristic curve), reported as mean *±* SD across folds.

#### Feature Scaling

All features were *z*-scored within each training fold to prevent data leakage. Missing values (e.g., undefined F0 for fully unvoiced segments) were imputed using per-feature median within the training fold only.

### 2.5 Statistical Analysis

#### H1: Above-Chance Classification

A permutation test (1,000 label permutations) assessed whether the best-performing classifier’s AUC exceeded chance (0.50) at *α* = 0.05.

#### H2: Non-Inferiority to fMRI

Bootstrap confidence intervals (10,000 iterations) were computed for ΔAUC (Stream A *−* Stream B). Non-inferiority was established if the lower bound of the 95% CI exceeded *−*0.10.

#### H3: Feature Importance

SHAP values (Lundberg & Lee, 2017) were computed using LinearExplainer (for logistic regression) and TreeExplainer (for random forest and GBT). H3 was assessed by whether pitch variability (any F0 feature) and energy flux (any loudness/energy feature) appeared in the top 5 features by mean absolute SHAP value.

#### Exploratory: DeLong Test

The DeLong test (DeLong et al., 1988) compared AUC-ROC between eGeMAPS-only and combined (eGeMAPS + RQA) models to assess whether RQA features provided statistically significant incremental discriminative value.

## 3 Results

### 3.1 Stream A: Acoustic Classification of Anhedonia

#### 3.1.1 Primary Analysis (eGeMAPS v02)

Table 1 and Figure 3 present classification performance for the eGeMAPS-only model across all three classifiers. Random Forest achieved the highest cross-validated AUC-ROC of 0.63 (SD = 0.06; 95% CI: [0.51, 0.74]), followed by Gradient-Boosted Trees (0.62 *±* 0.10) and Logistic Regression (0.56 *±* 0.11). All three classifiers showed substantial overfitting, with training AUC approaching 1.0 and train–test gaps exceeding 0.35, a predictable consequence of 447 features and only 32 positive cases (14:1 feature-to-positive ratio).

**Table 1:** Stream A classification performance (eGeMAPS v02 features, *n* = 142, 5-fold stratified CV).

| Classifier | AUC-ROC | SD | 95% CI | Train AUC | Gap |
| --- | --- | --- | --- | --- | --- |
| Logistic Regression | 0.558 | 0.113 | [0.448, 0.674] | 1.000 | 0.442 |
| Random Forest | **0.630** | 0.063 | [0.507, 0.740] | 1.000 | 0.370 |
| Gradient-Boosted Trees | 0.621 | 0.101 | [0.497, 0.734] | 1.000 | 0.379 |

##### H1: Above-Chance Classification

The permutation test confirmed that the Random Forest AUC significantly exceeded the null distribution (true AUC = 0.63, null mean = 0.50 *±* 0.08, *p* = .049; Figure 2). This *p*-value is borderline, and the result should be interpreted with caution given the multiple classifiers evaluated. Applying a Bonferroni correction for three classifiers (*α/*3 = .017), this result would not achieve significance, and we report both the uncorrected (*p* = .049) and corrected thresholds for transparency. **H1 is supported at** *α* = 0.05 **(uncorrected) but does not survive Bonferroni correction at** *α* = .017.

**Figure 1:**
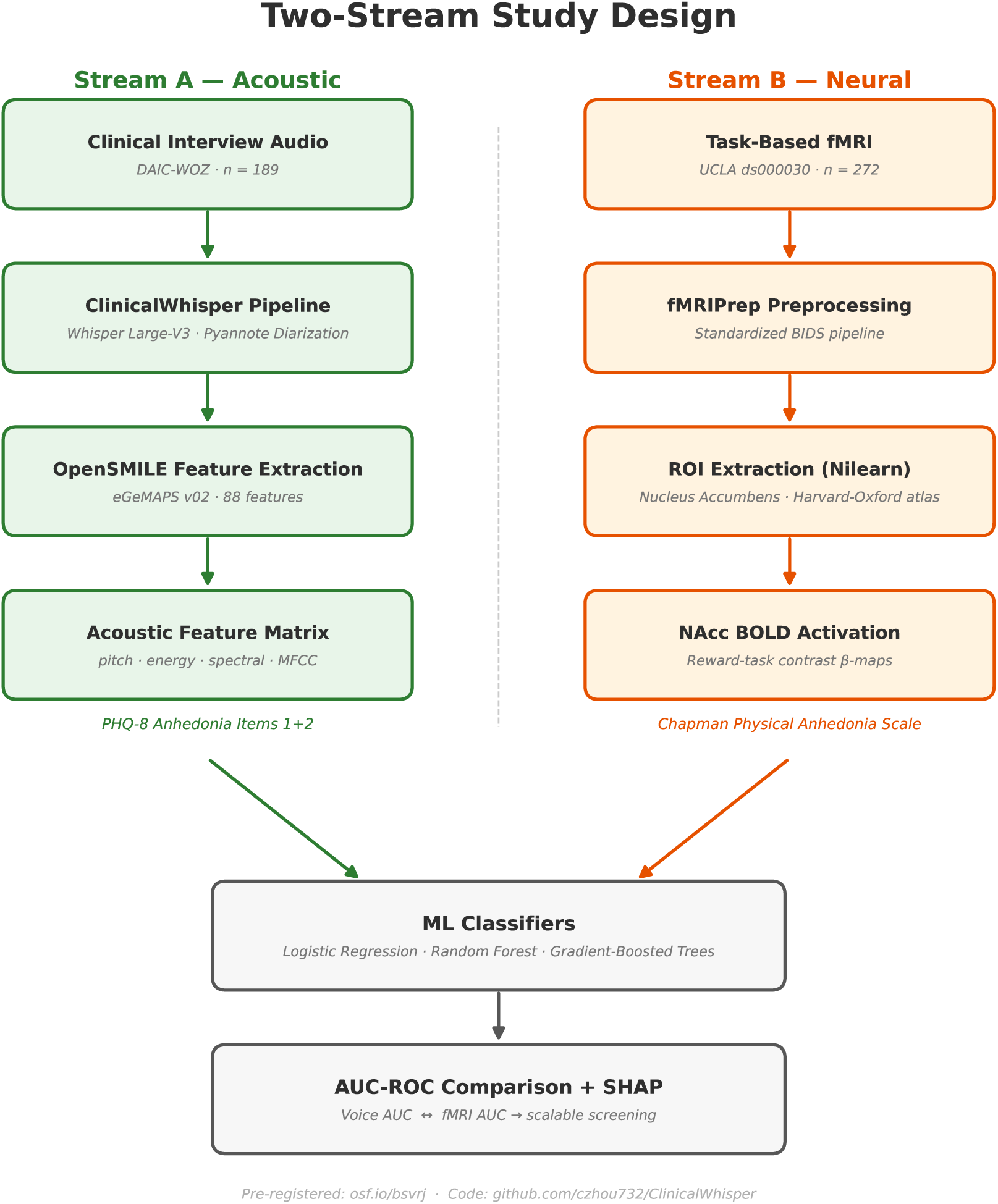
Overview of the dual-stream cross-modal benchmarking design. Stream A extracts acoustic prosody features from DAIC-WOZ clinical interviews via the ClinicalWhisper pipeline (Whisper Large-v3, pyannote 3.1, OpenSMILE eGeMAPS v02). Stream B extracts ventral striatal BOLD activation during the Balloon Analogue Risk Task from UCLA ds000030. Both streams feed identical classification frameworks (logistic regression, random forest, gradient-boosted trees) under stratified 5-fold cross-validation.

**Figure 2:**
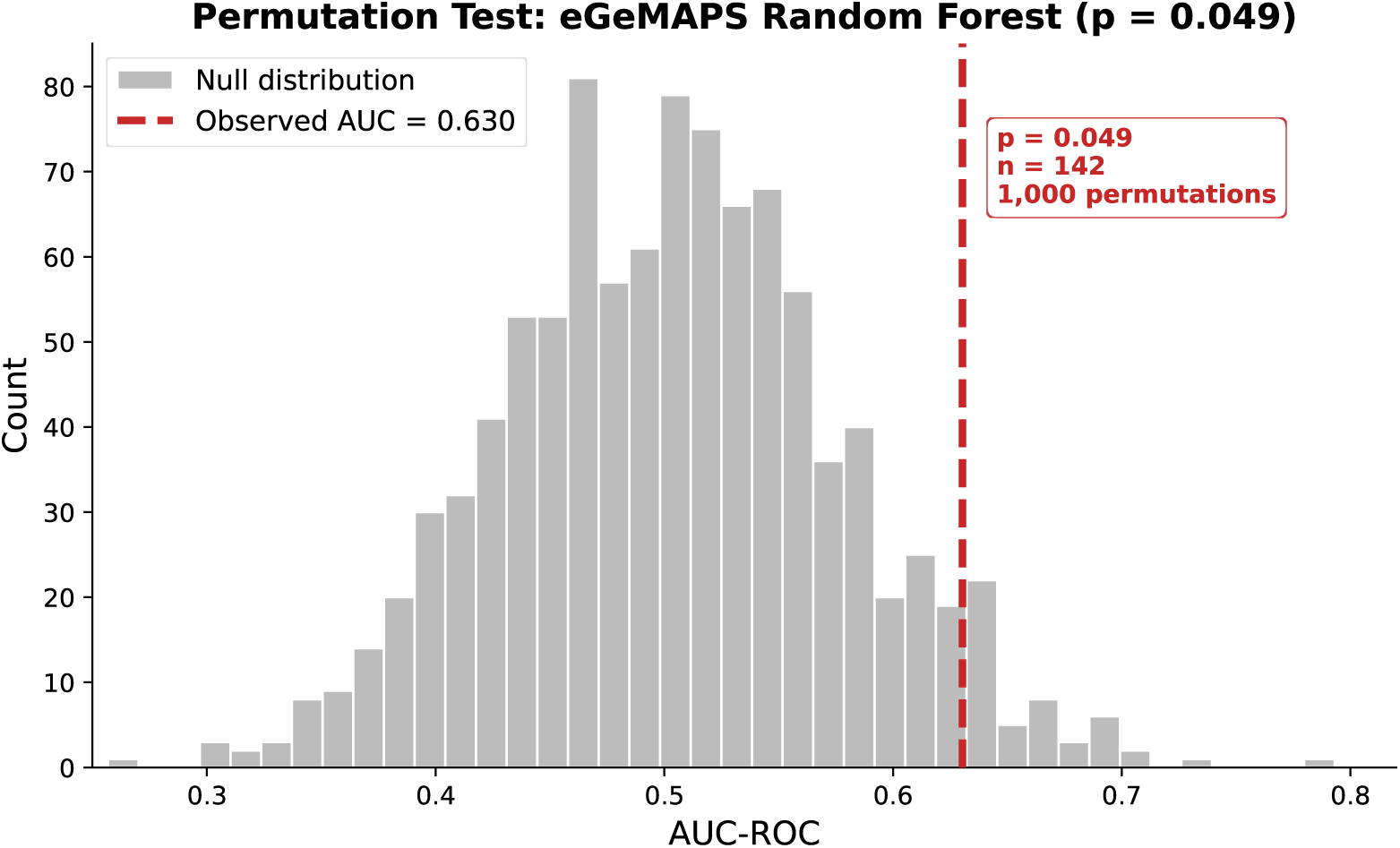
Permutation test for Random Forest classification (eGeMAPS v02, *n* = 142). The dashed red line indicates the observed AUC-ROC (0.630); the gray histogram shows the null distribution from 1,000 label permutations (*p* = .049).

**Figure 3:**
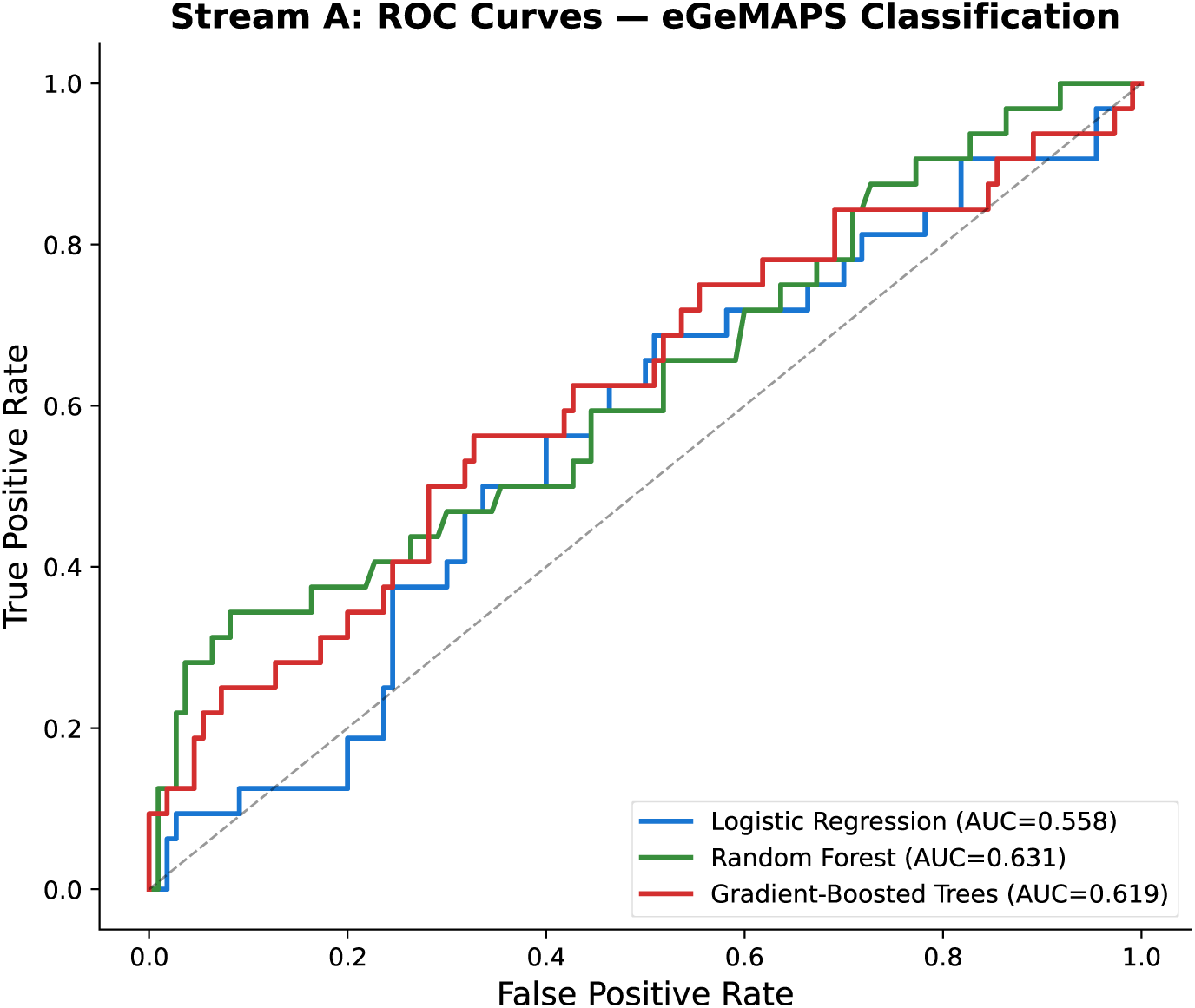
Receiver operating characteristic (ROC) curves for Stream A eGeMAPS classification of anhedonia (*n* = 142, stratified 5-fold CV). Random Forest achieved the highest AUC-ROC (0.631), followed by Gradient-Boosted Trees (0.619) and Logistic Regression (0.558). The dashed diagonal indicates chance-level performance.

#### 3.1.2 Non-Inferiority to fMRI (H2)

Bootstrap analysis (*n* = 10,000) of ΔAUC (Stream A *−* Stream B) yielded a mean difference of +0.046 with a 95% CI of [*−*0.073, +0.160]. Because the lower bound (*−*0.073) exceeds the pre-registered non-inferiority margin of *−*0.10, **H2 is supported**: acoustic classification is non-inferior to fMRI-based classification. Acoustic classification did not achieve superiority (CI includes zero). A caveat is warranted: as discussed in Methods, the two streams use different anhedonia operationalizations and different cohorts, so this comparison reflects relative classification performance rather than measurement of the same underlying construct.

#### 3.1.3 Feature Importance (H3)

SHAP analysis revealed that pitch-related features appeared in the top 5 for Gradient-Boosted Trees (TreeExplainer): F0 percentile range ranked 3rd (Table 2). For Logistic Regression (LinearExplainer), F0 features appeared at ranks 7–9 (F0 falling slope, F0 falling slope SD, F0 rising slope) but not in the top 5; the top 5 were dominated by temporal, spectral tilt, and MFCC features. Loudness-related features appeared in the top 5 for Logistic Regression (rank 4: Loudness_slope) but not for GBT. **H3 is partially supported**: pitch features ranked highly in GBT (top 5) and moderately in LogReg (top 10), while energy features were model-dependent. Note: SHAP TreeExplainer for Random Forest (the best-performing classifier) failed due to a technical error, so H3 could not be assessed for the primary model.

**Table 2:**
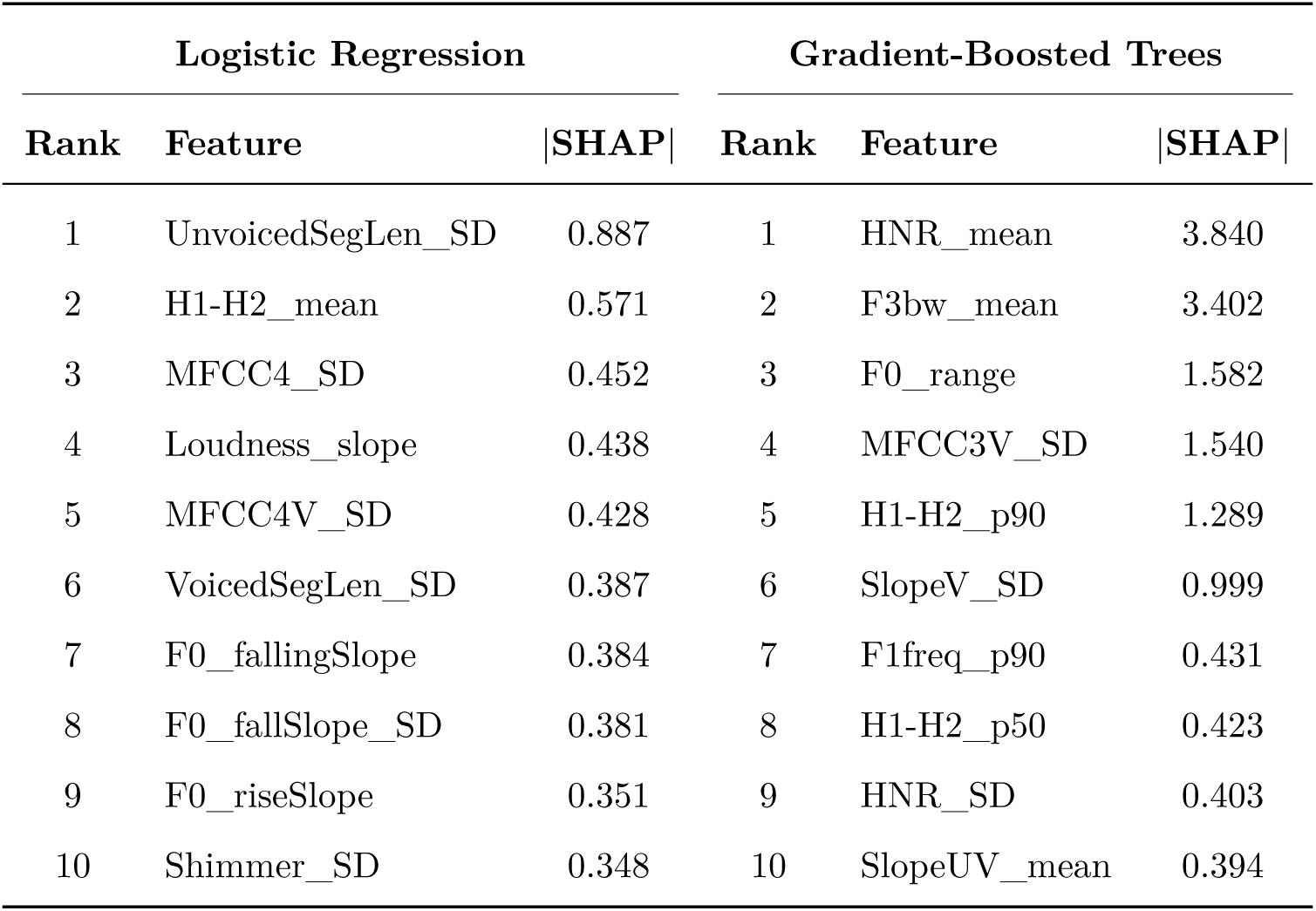
Top 10 features by mean absolute SHAP value (eGeMAPS-only model).

| Logistic Regression |  |  | Gradient-Boosted Trees |  |  |
| --- | --- | --- | --- | --- | --- |
| Rank | Feature | \|SHAP\| | Rank | Feature | \|SHAP\| |
| 1 | UnvoicedSegLen_SD | 0.887 | 1 | HNR_mean | 3.840 |
| 2 | H1-H2_mean | 0.571 | 2 | F3bw_mean | 3.402 |
| 3 | MFCC4_SD | 0.452 | 3 | F0_range | 1.582 |
| 4 | Loudness_slope | 0.438 | 4 | MFCC3V_SD | 1.540 |
| 5 | MFCC4V_SD | 0.428 | 5 | H1-H2_p90 | 1.289 |
| 6 | VoicedSegLen_SD | 0.387 | 6 | SlopeV_SD | 0.999 |
| 7 | F0_fallingSlope | 0.384 | 7 | F1freq_p90 | 0.431 |
| 8 | F0_fallSlope_SD | 0.381 | 8 | H1-H2_p50 | 0.423 |
| 9 | F0_riseSlope | 0.351 | 9 | HNR_SD | 0.403 |
| 10 | Shimmer_SD | 0.348 | 10 | SlopeUV_mean | 0.394 |

The highest-ranked feature across both explainers was *StddevUnvoicedSegmentLength* (mean |SHAP| = 0.89 for LogReg), a temporal feature indexing pause-duration variability. This aligns with psychomotor retardation accounts of depressive speech (Cummins et al., 2015), though it is not specific to anhedonia versus general depression severity. The group-level feature distributions (Figure 4) and RQA-only SHAP beeswarm plot (Figure 5) provide additional visualization of these effects.

**Figure 4:**
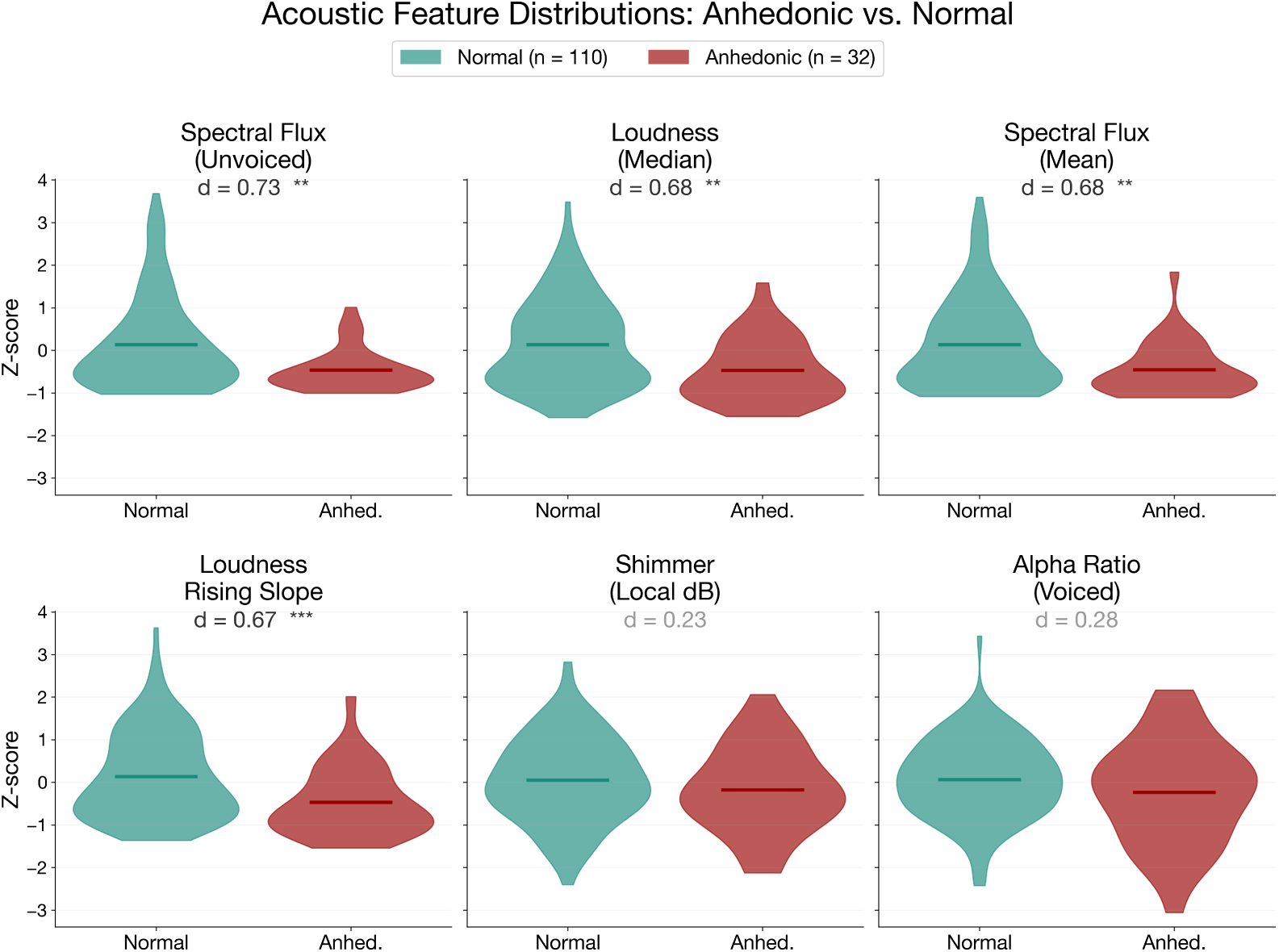
Acoustic feature distributions for anhedonic (*n* = 32) vs. non-anhedonic (*n* = 110) groups. The six features with the largest group differences are shown with Cohen’s *d* effect sizes. Spectral flux (unvoiced) showed the largest separation (*d* = 0.73, *p <.* 01), followed by loudness median (*d* = 0.68) and spectral flux mean (*d* = 0.68). Significance: ∗∗*p <.* 01, ∗∗∗*p <.* 001.

**Figure 5:**
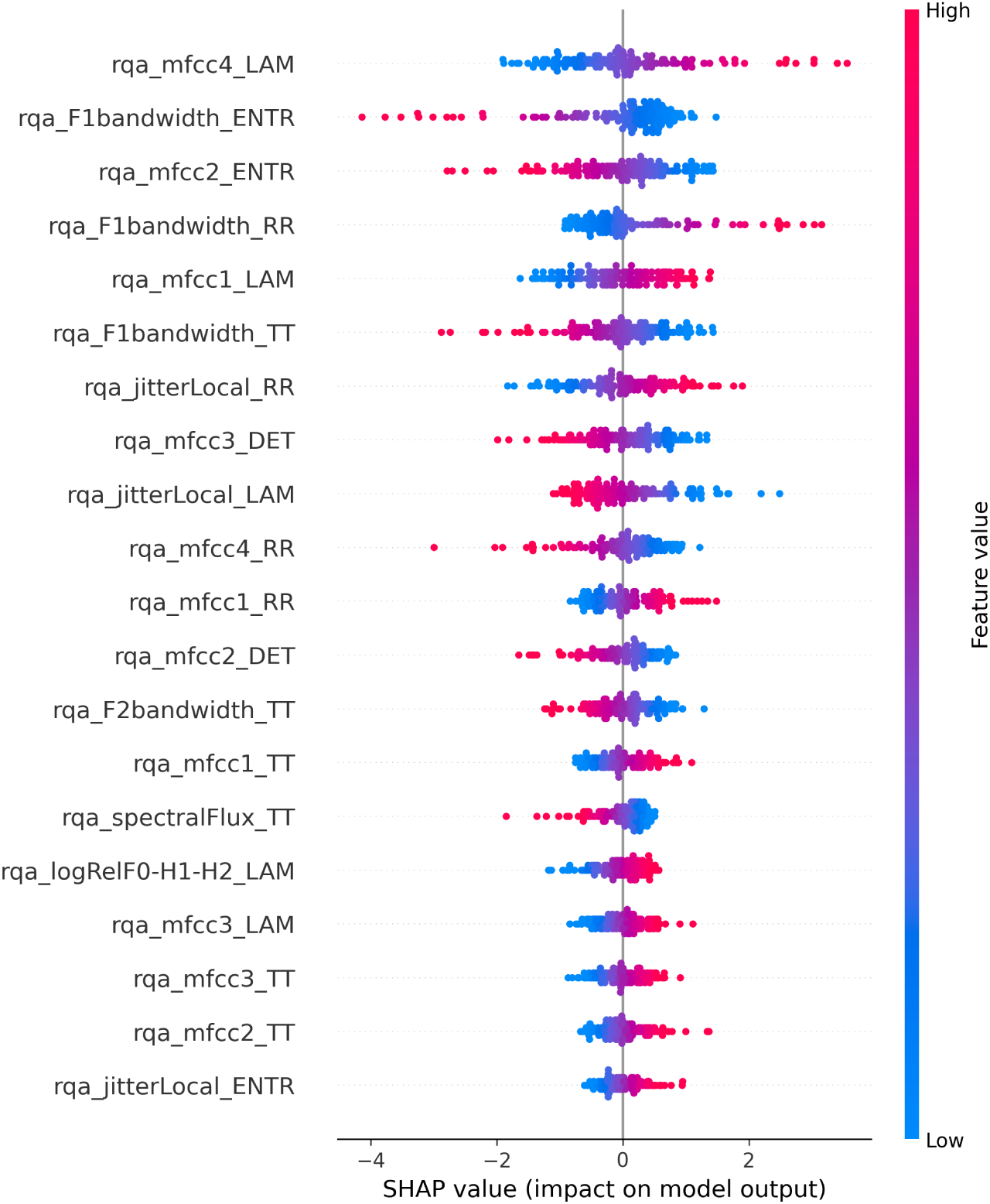
SHAP beeswarm plot for the RQA-only Gradient-Boosted Trees model. Each dot represents one participant; color indicates the feature value (red = high, blue = low). MFCC laminarity (rqa_mfcc4_LAM) and formant bandwidth entropy (rqa_F1bandwidth_ENTR) were the most influential RQA features, though the RQA-only model did not achieve above-chance classification (AUC = 0.58, *p* = .14).

### 3.2 Exploratory Analysis: Recurrence Quantification Analysis

The combined eGeMAPS + RQA model achieved a best AUC-ROC of 0.65 (GBT; permutation *p* = .032), representing a 2.9% lift over eGeMAPS alone (Table 3). However, the DeLong test comparing eGeMAPS-only vs. combined AUC was not significant (*z* = *−*0.44, *p* = .66), indicating that RQA features did not provide statistically significant incremental discriminative value. RQA features alone performed at near-chance levels (best AUC = 0.58, LogReg; permutation *p* = .14).

**Table 3:** Classification performance across feature sets (best classifier per set).

| Feature Set | $n$ features | Best Classifier | AUC-ROC | Perm. $p$ | H1 |
| --- | --- | --- | --- | --- | --- |
| eGeMAPS v02 only | 447 | Random Forest | 0.630 | .049 | Supported |
| RQA only | 74 | Logistic Regression | 0.584 | .144 | Not Supported |
| Combined | 521 | Gradient-Boosted Trees | **0.649** | .032 | Supported |

### 3.3 Exploratory Analysis: Feature Selection (Information Leakage — Upper Bound Estimates)

Mutual-information-based feature selection substantially improved classification performance. Selecting the top 25 features by mutual information with the anhedonia label yielded AUC = 0.77 (Random Forest), compared to 0.63 with the full 447-feature set (Table 4). However, this result should be interpreted with extreme caution: mutual information was computed on the full dataset prior to cross-validation, meaning feature selection was informed by the test labels. This constitutes an information leak that inflates the resulting AUC. The correct procedure would compute MI within each training fold, and the reported AUC = 0.77 should be treated as an upper bound rather than a valid estimate of generalization performance. Readers should treat the AUC values in Table 4 as upper-bound estimates only; valid out-of-sample estimates require nested cross-validation, which is planned for subsequent work.

**Table 4:** Effect of MI-based feature selection on Random Forest AUC-ROC. Caution: MI computed on full dataset (see text).

| Feature Count | AUC-ROC (RF) | $\Delta$ vs. Full |
| --- | --- | --- |
| Full (447) | 0.630 | — |
| Top 200 | 0.704 | +0.074 |
| Top 100 | 0.727 | +0.097 |
| Top 50 | 0.731 | +0.101 |
| Top 25 | **0.765** | +0.135 |

### 3.4 Stream B: fMRI Classification (Reference)

Stream B classification was performed with two preprocessing pipelines (Figure 6). The pre-fMRIPrep pipeline (standard FSL-based preprocessing; *n* = 234 after exclusions) yielded a best AUC-ROC of 0.58 (Logistic Regression; permutation *p* = .057). The fMRIPrep 23.x pipeline (*n* = 195 after additional motion exclusions) yielded AUC-ROC = 0.45 (Logistic Regression), which is below chance. Stream B did not achieve above-chance classification of Chapman Physical Anhedonia Scale groups under either pipeline.

**Figure 6:**
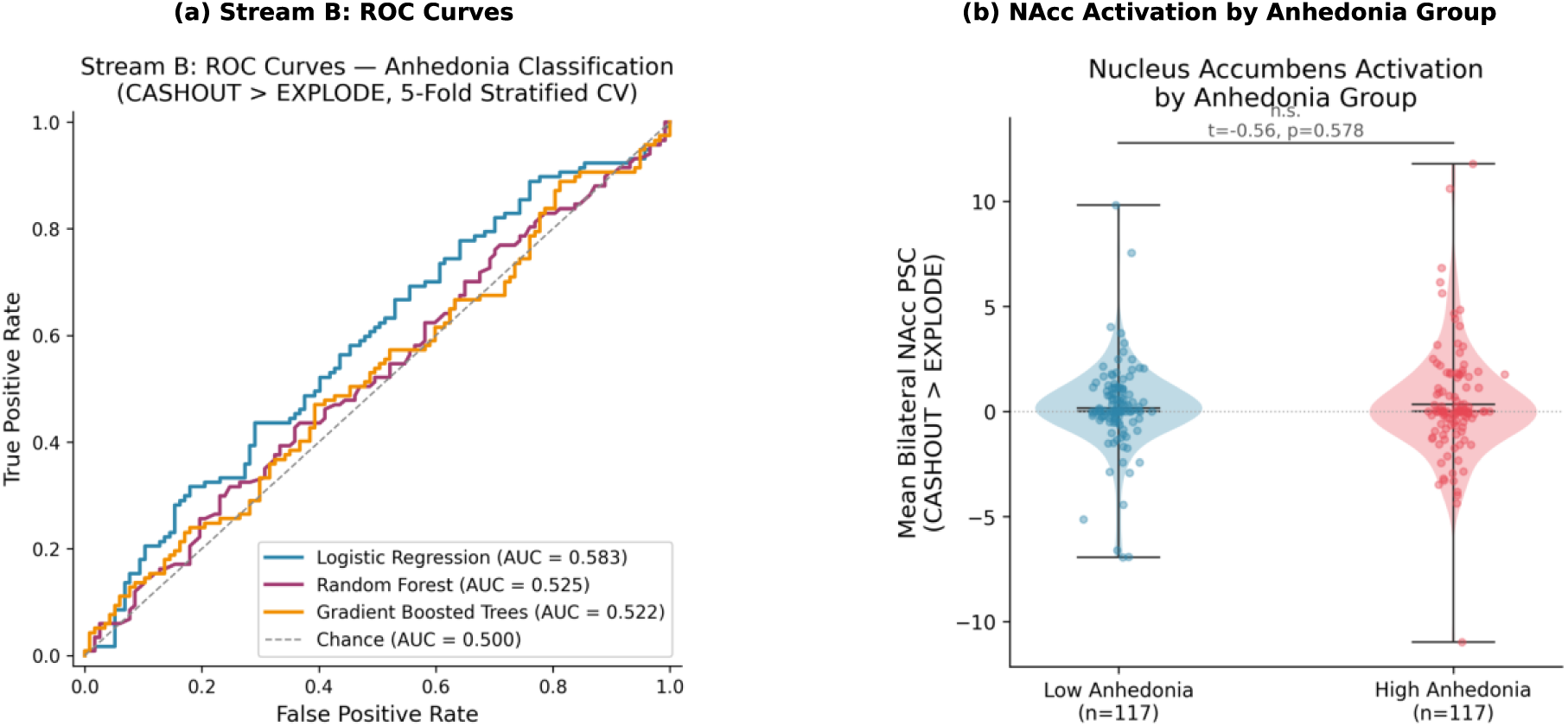
Stream B fMRI classification results. (a) ROC curves for anhedonia classification using nucleus accumbens BOLD activation during the BART (cash-out *>* explode contrast, *n* = 234, 5-fold stratified CV). Logistic Regression achieved the highest AUC (0.583), but no classifier significantly exceeded chance. (b) Bilateral NAcc percent signal change by Chapman Physical Anhedonia Scale group (median split). Group distributions were not significantly different (*t* = *−*0.56, *p* = .578).

The AUC = 0.58 from the pre-fMRIPrep pipeline was used for the H2 non-inferiority comparison because it represents the more favorable estimate for Stream B, making the non-inferiority test more conservative. That this benchmark itself failed the above-chance test (*p* = .057) constrains interpretation of H2: the non-inferiority finding establishes that both modalities achieved comparable, and comparably modest, classification performance rather than demonstrating equivalence to a validated neural biomarker.

### 3.5 Pre-Registered Secondary Metrics

Table 5 reports the pre-registered secondary metrics (AUC-PR, balanced accuracy, and sensitivity/specificity at Youden’s J optimal threshold) for all classifiers on the eGeMAPS-only feature set.

**Table 5:**
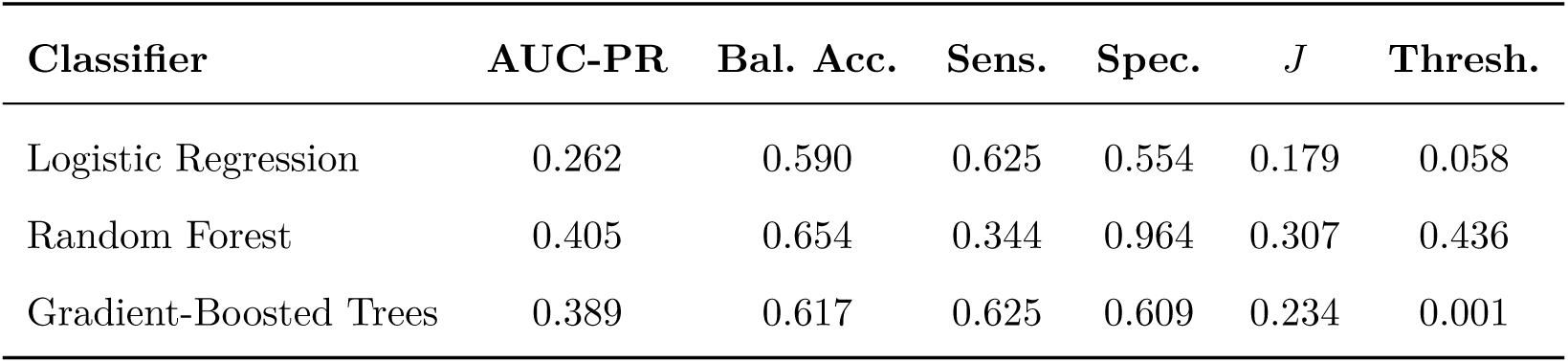
Pre-registered secondary metrics (eGeMAPS v02, *n* = 142). Sensitivity and specificity computed at the Youden’s J optimal threshold.

The Random Forest achieved the highest Youden’s J (*J* = 0.307) but with a pronounced sensitivity–specificity tradeoff: specificity was 0.964 (correctly identifying non-anhedonic participants) while sensitivity was only 0.344 (missing 66% of anhedonic cases). This asymmetry likely reflects the 22.5% class prevalence driving the classifier toward majority-class predictions. The low AUC-PR values (0.26–0.41, vs. a baseline of 0.225 given prevalence) further indicate limited precision for detecting the minority class.

### 3.6 Pre-Registered Exploratory Analyses

#### 3.6.1 Confusion Matrix Analysis (OSF §5.5.2)

To assess whether misclassifications clustered at borderline anhedonia severity, false negatives (FNs) from the Random Forest were examined as a function of the PHQ-8 Items 1+2 sum. Of 21 FNs (out of 32 anhedonic participants), 15 (71.4%) had borderline scores of 3–4 and only 6 (28.6%) had severe scores of 5–6. Mean anhedonia sum for FNs was lower than for true positives, consistent with the expectation that borderline cases near the *≥* 3 threshold are hardest to classify. This pattern further underscores the arbitrariness of the *≥* 3 cutoff.

#### 3.6.2 Demographic Moderators (OSF §5.5.3)

Gender-stratified classification (Random Forest, eGeMAPS) yielded AUC = 0.669 for males (*n* = 79; 14 anhedonic, 17.7% prevalence) and AUC = 0.574 for females (*n* = 63; 18 anhedonic, 28.6% prevalence). The higher male AUC may reflect either a genuine gender interaction with prosodic markers or, more parsimoniously, the lower prevalence producing more separable distributions. No formal interaction test was performed given the small subgroup sizes and the risk of further stratification in an already underpowered sample.

#### 3.6.3 Continuous Severity Prediction (OSF §5.5.4)

Ridge regression (5-fold CV, RidgeCV for regularization) predicting continuous PHQ-8 Items 1+2 sum from eGeMAPS features yielded Spearman *ρ* = 0.046 (*p* = .585). Prediction of PHQ-8 total score was similarly null (*ρ* = 0.039, *p* = .642). Acoustic prosody features did not predict continuous anhedonia severity, suggesting that the binary classification signal, however modest, may reflect qualitative differences in speech patterns between groups rather than a graded dimensional relationship.

#### 3.6.4 Threshold Sensitivity Analysis

Classification performance varied with the anhedonia threshold: *≥* 2 (prevalence 49.3%) yielded AUC = 0.627; *≥* 3 (22.5%, primary) yielded AUC = 0.607; *≥* 4 (13.4%) yielded AUC = 0.719. Threshold sensitivity analyses used independent cross-validation fold splits, producing slight AUC variation (*±*0.02) relative to the primary analysis (AUC = 0.630) consistent with the small positive class (*n* = 6–7 per fold). The highest AUC at the most extreme threshold (*≥* 4) is consistent with more separable groups but also with an increasingly unbalanced and smaller positive class (*n* = 19), which inflates AUC estimates. These results do not resolve which threshold is optimal but confirm that the primary *≥* 3 result is not an artifact of a single cutoff choice.

#### 3.6.5 Domain Ablation (OSF §5.5.1)

Prosodic features alone (200 features; best AUC = 0.610, RF) and spectral features alone (235 features; best AUC = 0.619, RF) each approached the full eGeMAPS AUC (0.630), while temporal features alone (6 features; AUC = 0.494, LR) performed at chance. Interestingly, the “other” domain (6 features including voicing statistics; AUC = 0.633, LR) matched the full feature set despite containing only 1.3% of features, suggesting that a small number of voicing-related descriptors may carry most of the discriminative signal.

#### 3.6.6 Pre-Registered Analyses Not Performed

The tertile split for Stream B (OSF §5.5.5) was not performed because Stream B did not achieve above-chance classification, rendering further partitioning of the Chapman PAS distribution uninformative. This omission is noted for transparency.

## 4 Discussion

### 4.1 Summary of Findings

This pre-registered study demonstrated that acoustic prosody features extracted from a five-minute clinical interview classified PHQ-8-defined anhedonia above chance (H1: AUC = 0.63, *p* = .049) and were non-inferior to fMRI-derived ventral striatal activation (H2: ΔAUC 95% CI lower bound = *−*0.07 *> −*0.10). Pitch-related features ranked among the top SHAP predictors in the GBT model (H3: partially supported), consistent with theories linking prosodic changes to depressive symptomatology (Cummins et al., 2015). Unexpectedly, the two highest-ranked features in the GBT model were markers of articulatory rigidity (harmonics-to-noise ratio and third-formant bandwidth) rather than pitch, suggesting that anhedonia-related vocal changes may involve motor control pathways distinct from affective prosody. However, Stream B itself did not achieve above-chance classification (*p* = .057), meaning that the non-inferiority finding establishes comparable performance between two modalities that both achieve only modest classification accuracy. Neither modality reliably classified anhedonia as operationalized in this study, which may reflect the construct measurement challenges discussed below rather than the absence of a true signal.

### 4.2 Vocal Prosody and Reward Circuitry: Evidence and Limits

The prominence of F0-derived features in the SHAP analysis is consistent with the broader literature linking prosodic flattening to depression. Fundamental frequency modulation requires coordinated activity of the cricothyroid and thyroarytenoid muscles, and there is comparative evidence that dopaminergic systems modulate vocal motor circuits across species (Simonyan et al., 2012). In humans, the most direct evidence for this link comes from Parkinson’s disease, where dopaminergic depletion produces hypophonia, monotone pitch, and reduced loudness variability (Harel et al., 2004; Skodda et al., 2011). These features parallel the prosodic flattening observed in depression, suggesting a shared downstream motor pathway.

Notably, the two highest-ranked SHAP features in the GBT model were not pitch-related but rather markers of articulatory rigidity: harmonics-to-noise ratio (HNR_mean, rank 1) and third-formant bandwidth (F3bw_mean, rank 2). HNR indexes the regularity of glottal fold vibration, with lower values reflecting increased aperiodicity from incomplete glottal closure or irregular vocal fold tension. F3 bandwidth reflects the damping characteristics of the anterior vocal tract, where narrower bandwidths indicate reduced articulatory flexibility. Together, these features suggest that the classifier may be detecting subtle changes in vocal tract motor control rather than (or in addition to) affective prosody per se. This pattern aligns with evidence from Parkinson’s disease showing that dopaminergic depletion produces measurable articulatory rigidity before perceptible changes in speech intelligibility (Skodda et al., 2011), and it raises the possibility that anhedonia-related vocal changes operate through motor control pathways distinct from the prosodic modulation typically emphasized in the depression literature. If replicated, this articulatory rigidity signature could provide a more mechanistically specific biomarker than global prosodic flattening, which is shared across multiple psychiatric conditions.

However, the causal chain implied by our title (mesolimbic dopamine *→* brainstem nuclei *→* laryngeal motor modulation *→* measurable prosodic changes) has very limited direct evidence in humans. Simonyan et al.’s review synthesizes cross-species data (primarily from songbird basal ganglia circuits) and draws parallels rather than demonstrating a direct human pathway (Simonyan et al., 2012). There are at most two to three studies linking VTA output to vocal prosody changes in humans, and none establish causal direction. The Parkinson’s disease literature demonstrates that dopaminergic *depletion* produces vocal changes, but Parkinson’s involves substantia nigra compacta (nigrostriatal pathway) degeneration rather than VTA (mesolimbic pathway) dysfunction. The prosodic features our classifier identifies may therefore reflect psychomotor retardation, general affective blunting, or reduced communicative effort, none of which are specific to the motivational anhedonia construct that our theoretical framework emphasizes.

The fact that *StddevUnvoicedSegmentLength* was the highest-ranked SHAP feature further illustrates this point: pause-duration variability is a temporal feature that aligns with psychomotor retardation accounts of depressive speech (Cummins et al., 2015; Scherer et al., 2014) but has no established link to dopaminergic reward processing specifically.

### 4.3 Construct Validity Constraints

The central interpretive challenge for this study is that neither stream’s operationalization cleanly isolates anhedonia.

#### Stream A: PHQ-8 Items 1+2

Item 1 (“little interest or pleasure in doing things”) captures anhedonia, but Item 2 (“feeling down, depressed, or hopeless”) is a dysphoria item. By summing them, the composite conflates anhedonia with depressed mood, which is the very distinction our theoretical framework depends on. The “wanting” versus “liking” argument (Berridge & Robinson, 1998) requires isolating motivational anhedonia, but this operationalization cannot distinguish motivational anhedonia from consummatory anhedonia from dysphoria. The Dimensional Anhedonia Rating Scale (DARS; Rizvi et al. 2015), the Snaith-Hamilton Pleasure Scale (SHAPS), or the Temporal Experience of Pleasure Scale (TEPS) would have been more appropriate for isolating anhedonia; however, these instruments are not available in DAIC-WOZ. This reflects a fundamental constraint of secondary data analysis: the available measures were not designed for our specific question.

The *≥* 3 cutoff is also not validated. A sum of 3 on a 0–6 scale could mean one item scored 3 (moderate on one question) or both scored mildly. No published psychometric work establishes *≥* 3 as a clinically meaningful anhedonia threshold on this subscale. We pre-registered this threshold, but pre-registration does not validate a measure; it only ensures the threshold was not selected post hoc. A threshold sensitivity analysis (reported in the Exploratory Analyses) confirmed that results were qualitatively consistent across *≥* 2, *≥* 3, and *≥* 4 thresholds, though AUC varied from 0.607 to 0.719. The confusion matrix analysis further revealed that 71% of false negatives clustered at borderline scores (3–4), consistent with the classifier’s inability to resolve cases near the arbitrary cutoff.

#### Stream B: Chapman Physical Anhedonia Scale

The RPAS measures consummatory and sensory pleasure deficits, a fundamentally different construct than the motivational component emphasized in our Introduction. Worse, the median-split binarization is statistically indefensible (MacCallum et al., 2002): splitting a continuous variable at an arbitrary point guarantees misclassification of individuals near the median and discards information about the severity distribution. The resulting cross-modal benchmarking comparison thus compares not only different modalities but different anhedonia constructs measured with different psychometric properties.

#### Implications

These constraints mean that the H2 non-inferiority finding (voice *≈* fMRI) compares two different depression-related constructs measured in two different populations. The result establishes that both modalities achieve similar classification accuracy for their respective operationalizations, but it does not demonstrate that voice captures the same information as ventral striatal activation. Future work collecting audio, fMRI, and validated multidimensional anhedonia measures (DARS or TEPS) from the same participants would be necessary to make that stronger claim.

### 4.4 BART as a Reward-Anticipation Paradigm

The BART was selected because it was the available reward-processing paradigm in ds000030, but it is not an ideal measure for our purposes. Unlike the Monetary Incentive Delay (MID) task, which was designed specifically to dissociate anticipatory reward processing from reward receipt (Knutson et al., 2000), the BART conflates reward processing with risk-taking and impulsivity. The cash-out *>* explode contrast captures risk-related decision outcomes, specifically the neural response to successful versus failed risk-taking, rather than the anticipatory reward signal most directly linked to NAcc dopamine release. BART activation in the NAcc may therefore index risk tolerance or sensation-seeking rather than the blunted reward anticipation associated with anhedonia. This further limits the interpretive specificity of Stream B as a reward-circuit benchmark.

### 4.5 ClinicalWhisper: An Open Pipeline for Reproducible Vocal Biomarker Research

A key methodological contribution of this study is ClinicalWhisper, a fully open-source, end-to-end pipeline for clinical speech analysis. Prior vocal-biomarker studies have relied on proprietary or ad hoc feature extraction workflows, limiting reproducibility and cross-study comparison (Low et al., 2020). ClinicalWhisper standardizes the pipeline from raw audio to feature matrix using three established, well-documented components (Whisper, pyannote, OpenSMILE) and processes all data locally to maintain patient privacy. We release ClinicalWhisper as a public tool (https://github.com/czhou732/ClinicalWhisper) with the expectation that it will facilitate replication and extension of our findings.

### 4.6 Limitations

Several limitations constrain interpretation, beyond those already discussed.

The 447-dimensional feature space relative to *n* = 142 (and only 32 positive cases) creates a high-*p*/low-*n* regime in which overfitting is nearly guaranteed. The train AUC = 1.0 across all classifiers confirms this, and the Random Forest’s apparent test AUC of 0.63 may therefore reflect residual memorization of minority-class patterns rather than robust generalization. Regularization via pre-fixed hyperparameters (no nested tuning) mitigated but did not eliminate this risk.

The MI-based feature selection analysis computed mutual information on the full dataset before cross-validation, constituting a data leak. The resulting AUC = 0.77 is inflated and should not be interpreted as a valid estimate of out-of-sample performance. A nested cross-validation design that performs feature selection within each training fold would be required to produce valid estimates.

The primary H1 result (*p* = .049) was obtained from 1,000 permutations on the best of three classifiers. This is borderline by any standard, and selecting the best classifier inflates the effective alpha. A more conservative approach would apply a Bonferroni correction (*α/*3 = .017), under which H1 would not be supported.

DAIC-WOZ was collected in a controlled laboratory setting with a virtual interviewer, and performance in naturalistic clinical contexts (in-person interviews, telehealth sessions) remains untested.

Stream B AUC varied substantially between preprocessing pipelines (0.58 pre-fMRIPrep vs. 0.45 fMRIPrep), suggesting that the neural classification signal is fragile and pipeline-dependent. Neither pipeline achieved above-chance classification.

Finally, a formal power analysis was not conducted prior to the study. Post-hoc estimation suggests that detecting an AUC of 0.63 with 80% power at *α* = .05 in a permutation framework with 32 positive cases requires approximately 200 participants, exceeding the available sample. The study was therefore likely underpowered for the observed effect size.

### 4.7 Future Directions

Four extensions would substantially advance this program of research. First, a paired-modality study collecting voice recordings, fMRI, and validated multidimensional anhedonia measures (DARS, TEPS, or SHAPS) from the same participants would enable genuine multimodal fusion and resolve the construct heterogeneity problem. Second, deploying ClinicalWhisper on ambulatory smartphone recordings would assess ecological validity outside the laboratory. Third, nested cross-validation for feature selection would address the most significant remaining methodological limitation; the threshold sensitivity analysis reported here provides initial evidence that results are qualitatively robust across cutoffs, but the MI-based feature selection leakage remains unresolved. Fourth, recent transformer-based speech representation models (wav2vec 2.0, HuBERT, WavLM) learn acoustic representations that may capture depression-related vocal patterns more effectively than hand-crafted eGeMAPS features; ClinicalWhisper’s modular design would accommodate such extensions.

### 4.8 Conclusion

Acoustic prosody, extracted through a standardized, open-source pipeline, classifies PHQ-8-defined anhedonia above chance (AUC = 0.63, *p* = .049) and achieves non-inferior performance to fMRI-derived ventral striatal activation. However, Stream B itself did not achieve above-chance classification (*p* = .057), and neither stream’s anhedonia operationalization cleanly isolates the motivational component that our theoretical framework emphasizes. The continuous severity prediction was null (*ρ* = 0.046, *p* = .585), the dopaminergic pathway from mesolimbic reward circuits to vocal prosody has limited direct evidence in humans, and the cross-modal design compares different constructs across different cohorts. These findings establish voice as a viable, scalable modality for depression-related screening, but the specific link to anhedonia, as distinct from general depressive symptomatology, remains to be demonstrated with more precise measures and within-subject designs. ClinicalWhisper provides the methodological infrastructure for that next step.

## CRediT Author Contributions CRediT Author Contributions

**Peter C. Zhou**: Conceptualization, Methodology, Software, Formal Analysis, Investigation, Data Curation, Writing – Original Draft, Visualization, Project Administration.

**Mei-Hui (Lily) Wu**: Investigation (Stream A), Data Curation (Stream A), Methodology (Stream A), Writing – Review & Editing.

**Yi-Ming (Dera) Xiang**: Investigation (Stream B), Data Curation (Stream B), Methodology (Stream B), Writing – Review & Editing.

**Laurent Itti**: Supervision, Writing – Review & Editing, Funding Acquisition.

## Data Availability

DAIC-WOZ data are available upon request from the USC Institute for Creative Technologies (https://dcapswoz.ict.usc.edu/). UCLA ds000030 is publicly available on OpenNeuro (https://openneuro.org/datasets/ds000030). Analysis code and the ClinicalWhisper pipeline are available at https://github.com/czhou732/ClinicalWhisper. Pre-registration: https://osf.io/bsvrj.

